# dadasnake, a Snakemake implementation of DADA2 to process amplicon sequencing data for microbial ecology

**DOI:** 10.1101/2020.05.17.095679

**Authors:** Christina Weiβbecker, Beatrix Schnabel, Anna Heintz-Buschart

**Affiliations:** Helmholtz Centre for Environmental Research GmbH - UFZ, Department of Soil Ecology; German Centre for Integrative Biodiversity Research (iDiv) Halle-Jena-Leipzig, Bioinformatics Unit

**Keywords:** rRNA gene sequence analysis, denoising, exact sequence variants, R, pipeline, microbiome, community structure

## Abstract

**Background:** Amplicon sequencing of phylogenetic marker genes, e.g. 16S, 18S or ITS rRNA sequences, is still the most commonly used method to determine the composition of microbial communities. Microbial ecologists often have expert knowledge on their biological question and data analysis in general, and most research institutes have computational infrastructures to employ the bioinformatics command line tools and workflows for amplicon sequencing analysis, but requirements of bioinformatics skills often limit the efficient and up-to-date use of computational resources.

**Results:** dadasnake wraps pre-processing of sequencing reads, delineation of exact sequence variants using the favorably benchmarked, widely-used the DADA2 algorithm, taxonomic classification and post-processing of the resultant tables, and hand-off in standard formats, into a user-friendly, one-command Snakemake pipeline. The suitability of the provided default configurations is demonstrated using mock-community data from bacteria and archaea, as well as fungi.

**Conclusions:** By use of Snakemake, dadasnake makes efficient use of high-performance computing infrastructures. Easy user configuration guarantees flexibility of all steps, including the processing of data from multiple sequencing platforms. dadasnake facilitates easy installation via conda environments. dadasnake is available at https://github.com/a-h-b/dadasnake.

## Findings

### Background

Since the first reports 15 years ago [1], high-throughput amplicon sequencing has become the most common approach to monitor microbial diversity in environmental samples. Sequencing preparation, throughput and precision have been consistently improved, while costs have decreased. Computational methods have been refined in the recent years, especially with the shift to exact sequencing variants and better use of sequence quality data [2,3]. While amplicon sequencing can have severe limitations, such as limited and uneven taxonomic resolution [4,5], over- and underestimation of diversity [6,7], lack of absolute abundances [8,9] and missing functional information, amplicon sequencing is still considered the method of choice to gain an overview of microbial diversity and composition in a large number of samples [10,11]. Consequently, the sizes of typical amplicon sequencing datasets have grown. In addition, synthesis efforts are undertaken, requiring efficient processing pipelines for amplicon sequencing data [12]. Due to the unique, microbiome-specific characteristics of each dataset and the need to integrate the community structure data with other data types, such as abiotic or biotic parameters, users of data processing tools need to have expert knowledge on their biological question and statistics. It is therefore desirable that workflows should be as user-friendly as possible. Several widely used tool collections e.g. QIIME 2 [13], mothur [14], usearch [15], vsearch [16], and one-stop pipelines, e.g. lOTUs [17], with new approaches continually being developed, e.g. OCToPUS [18], PEMA [19]. Typically, workflows balance learning curves, configurability and efficiency.

### Purpose of dadasnake

dadasnake is a workflow for amplicon sequencing data processing into annotated exact sequence variants. It is set up with microbial ecologists in mind, to be run on high-performance clusters without the users needing any expert knowledge on their operation. dadasnake is implemented in Snakemake [20] using the conda package management system. Consequently, it features a simple installation process, a one-command execution, and high configurability of all steps with sensible defaults. dadasnake includes example workflows for common applications and produces a unique set of useful outputs, comprising relative abundance tables with taxonomic and other annotations in multiple formats, reports on the data processing and visualizations of data quality at each step. The workflow is open-source, based on validated, favourably benchmarked tools.

### Implementation

The central processing within dadasnake wraps the DADA2 R package [21], which accurately determines sequence variants [22–24]. The dadasnake wrapper eases DADA2 use and deployment on computing clusters without the overhead of larger pipelines with DADA2 such as QIIME 2 [13]. Within dadasnake, the steps of quality filtering and trimming, error estimation, inference of sequence variants, and, optionally, chimera removal are performed (Figure 1). Prior to quality filtering, dadasnake optionally removes primers and re-orients reads using cutadapt [25]. Taxonomic classification is realized using the reliable naïve Bayes classifier as implemented in mothur [14], or by DECIPHER [26,27] with optional species-identification in DADA2. BLAST [28] can optionally be used to annotate all or only unclassified sequence variants. The sequence variants can be filtered based on length, taxonomic classification or recognizable regions, namely ITSx [29] before downstream analysis. For downstream analyses, a multiple alignment [30] and FastTree-generated tree [31] can be integrated into a phyloseq [32] object. Alternatively, tab-separated or R tables and standardized BIOM format (https://biom-format.org/index.html) are generated. dadasnake records statistics, including numbers of reads passing each step, quality summaries, error models, and rarefaction curves [33]. All intermediary steps and configuration settings are saved for reproducibility.

**Figure 1:**
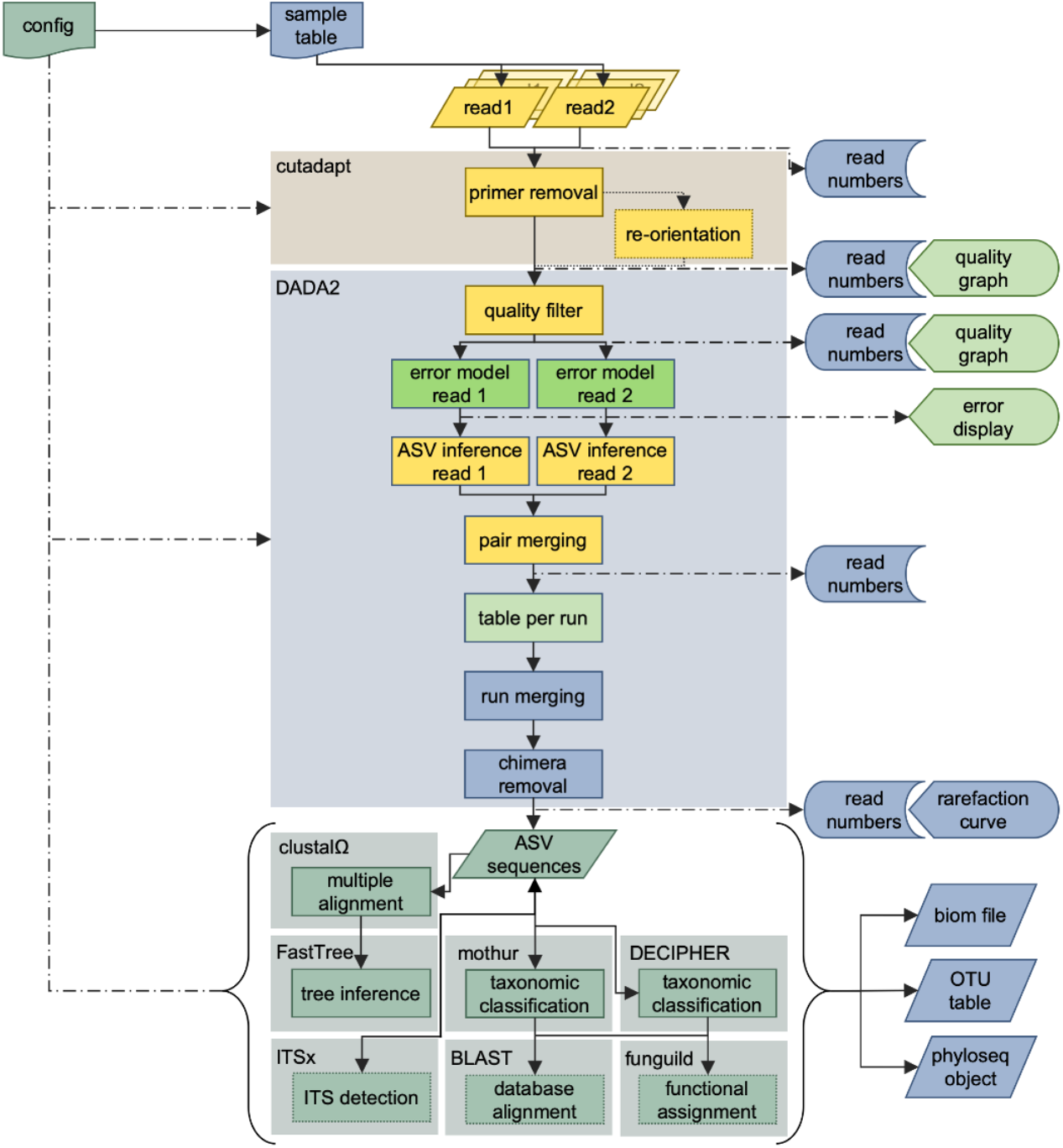
Overview of the dadasnake workflow for paired-end Illumina sequencing of a fungal ITS region with inputs (configuration file, sample table and read files) and outputs (read numbers, graphical representations of quality and error models, rarefaction curves and “OTU tables”, in biom, table and phyloseq format). The steps are configurable and alternative workflows exist, e.g. for single-end, non-Illumina datasets, or other target regions. Primer-removal and all post-DADA2-steps are optional. Colours represent the level of analysis: yellow – analysis per library/sample, bright green – analysis per run, sea green – analysis of the cumulated dataset; blue – analysis for the whole dataset with sample-wise documentation; note – the DADA2 block can be performed in pooled mode at the level of the whole dataset.

Reproducibility, user-friendliness, and modular design are facilitated by the Snakemake framework, a popular workflow manager for reproducible and scalable data analyses [20]. Snakemake also generates html reports, which store code, version numbers, the workflow and links to results. DADA2 and the other tools are packaged in conda environments to facilitate installation. For reasons of reproducibility, dadasnake uses fixed versions of all tools, which are regularly tested on mock-datasets and updated when improvements become available. Snakemake also ensures flexible use as single-threaded local workflow or efficient deployment on a batch scheduling system. Currently slurm and univa/sun grid engine scheduler configurations are defined for dadasnake.

### dadasnake configuration and execution

The whole dadasnake workflow is started with a single command (“dadasnake -c configuration.yaml”). The user provides a tab-separated table with sample names and input files, as well as a configuration file in the simple, human-readable and -writable YAML format (see Supplementary file 1 for a worked example) to determine which steps should be taken and with what settings (see description of all configurable parameters in Supplementary table 1). dadasnake is highly configurable compared to other Snakemake-based amplicon sequencing workflows, e.g. Hundo [34]. To facilitate its use, dadasnake provides easily adjustable, tested default settings and configuration files for several use cases.

dadasnake can use single end or paired end data. DADA2 can be efficiently employed by parallelizing most steps by processing samples individually (https://benjjneb.github.io/dada2/bigdata.html). Pooled analysis can alternatively be chosen in dadasnake and we recommend it for more error prone technologies such as 454 or third generation long reads. While DADA2 has been designed for Illumina technology [21], dadasnake has been tested on Roche pyrosequencing data [35] and circular consensus Pacbio [36] and Oxford Nanopore data [37,38] (see supporting material). dadasnake provides example configurations for these technologies and for Illumina-based analysis of 16S, ITS and 18S regions of bacterial and fungal communities.

dadasnake offers a range of different output formats for easy integration with downstream analysis tools. Tab-separated or R tables and standardized BIOM format (https://biom-format.org/index.html), or a phyloseq [32] object are generated as final outputs in the user-defined output directory (description of all outputs in Supplementary table 2). Visualizations of the input read quality, read quality after filtering, the DADA2 error models and rarefaction curves of the final dataset are also saved into a stats folder within the output. The numbers of reads passing each step are recorded for trouble shooting. All intermediary steps and configuration settings are saved for reproducibility and to restart the workflow in case of problematic settings or datasets. The Snakemake-generated html report contains all software versions and settings to facilitate the publication of the workflow’s results (see supporting material).

Snakemake provides detailed error reports and the logs of each step are recorded during runs. E-mail notifications of start and finishing can be sent. Users can find trouble shooting help and file issues at https://github.com/a-h-b/dadasnake.

### Use cases: performance

To demonstrate dadasnake’s performance, public datasets of different scales were processed. The performance of dadasnake depends strongly on the number of reads, number of samples, number of ASVs, and the required processing steps.

Small datasets can be run on single cores with less than 8 GB RAM, but profit from dadasnake’s parallelization. For example, a 24-sample dataset with 2.9 million 16S rRNA V4 reads [39] could be completely processed, including preprocessing, quality filtering, ASV determination, taxonomic assignment, treeing, visualization of quality, and hand-off in various formats with a total walltime of 150 minutes. Running time was reduced to 100 minutes, when four cores were used, especially due to the parallelization of the preprocessing and ASV determination steps (Fig. 2 a&b). Hardware requirements for small datasets are minimal, including small personal laptops. A medium-sized ITS1 dataset (267 samples with a total of 46.8 million reads [40]) could be processed in just under 4 hours on four 8 GB cores, including quality filtering, ASV determination, extraction of ITS1, taxonomic assignment, visualization of quality, and hand-off in various formats (Fig. 2 c). While the system walltime was similar, the use of 15 cores reduced the runtime by a factor 2 (Fig. 2 d).

**Figure 2:**
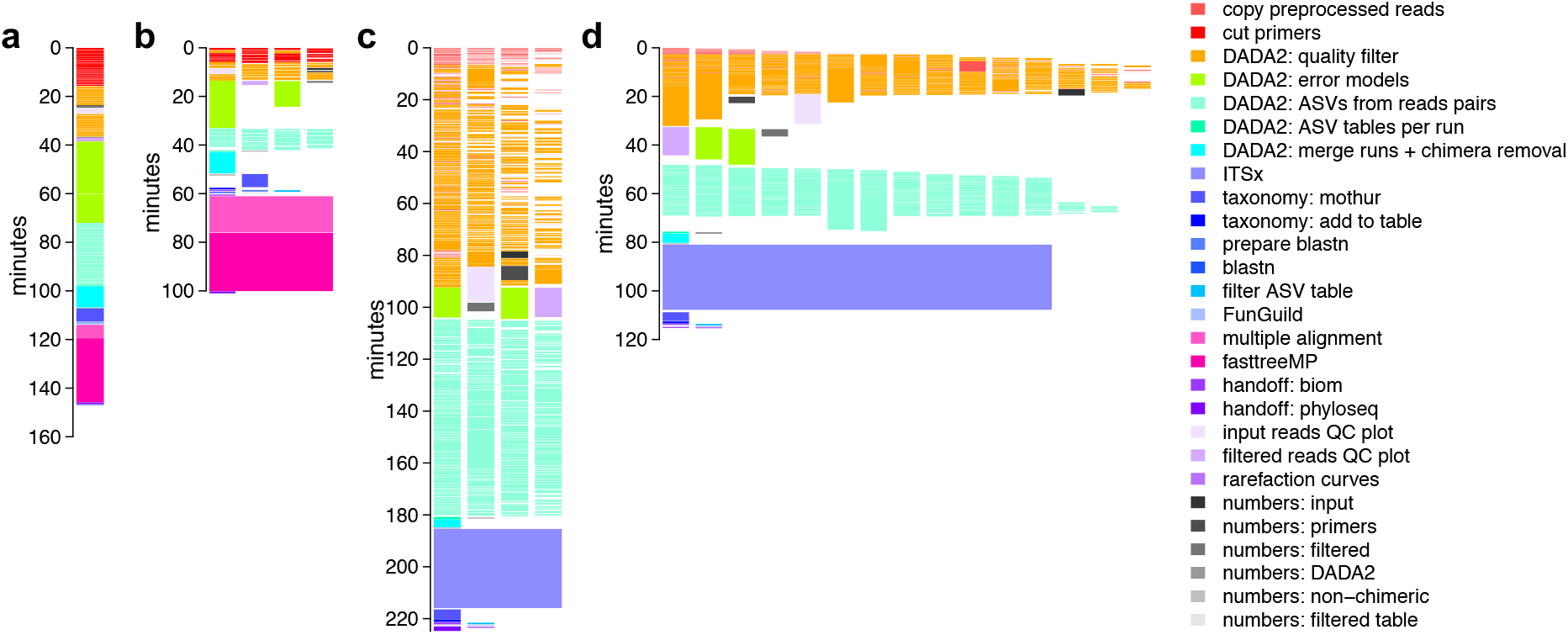
Visualization of resource use by processing different datasets. a) The small (24 sample) 16S rRNA V4 amplicon dataset *[39]* processed linearly on a single core; b) the same dataset processed on up to four cores (each depicted as a vertical stack); c) a medium sized (267 sample) ITS1 amplicon dataset *[40],* processed on up to four cores; d) the same dataset, processed on up to 15 cores. Each block represents one job issued by dadasnake, colors represent the respective steps.

Generally speaking, dadasnake’s parallelization of primer trimming, quality filtering, and ASV determination leads to shortened running times, while some steps, like merging of the ASV results of the single samples and all processing of assembled ASV tables, such as chimera removal, taxonomic annotation and treeing, are run sequentially. While dadasnake requests more cores for steps that use parallelized tools, such as ITSx or treeing, the speed-up is usually incremental. Of note for users of shared cluster environments, dadasnake does not occupy cores idly, e.g. when only a single core is used for merging of runs and chimera removal (Fig. 2 b-d) the other cores are available to other users, leading to high overall efficiency (> 90%).

dadasnake is able to preprocess reads, report quality, determine ASVs, and assign taxonomy for very large datasets, e.g. the original 2.1 billion reads in >27,000 samples of the Earth Microbiome Project publication [12] within 87 real hours on only up to 50 CPU cores. Due to the independent handling of the preprocessing, filtering and ASV definition steps, the number of input samples only prolongs the run time linearly. Sample merging and handling of the final table, however, requires more RAM the more unique ASVs and samples are found (e.g. > 190 GB for the >700,000 ASVs in the >27,000 samples of the Earth Microbiome Project). Tree building was not possible for this dataset on our infrastructure. For very large datasets it is therefore advisable to filter the final table before postprocessing steps.

### Use cases: accuracy

To demonstrate dadasnake’s potential to accurately determine community composition and richness, two mock community datasets from Illumina sequencing of bacterial and archaean [41] and fungal [42] DNA were analysed (compositions displayed in Supplementary table 3). In both cases, the genus-level composition was determined mostly correctly (Figure 2 a&b; supplementary table 3). One fungal taxon and two archaeal and three bacterial taxa were not detected at all, likely because they were not amplified. False positive bacterial genera were unrelated to the taxa in the mock-community and contained several human/skin-associated taxa, like *Corynebacterium* and *Staphylococcus,* as well as commonly detected sequencing contaminants like Rhizobiaceae and *Sphingomonas* (see overlap with [43] in Supplementary table 3). The large number of false-positives was therefore likely caused by contaminations in the bacterial dataset which have been observed in this dataset before [24]. For the fungal dataset, one *Fusarium* sequence was misclassified as *Giberella.* In the same settings, the ASV richness was inferred close to correctly at 59 and 19 prokaryotic and fungal ASVs, respectively (ignoring the contaminants; Figure 2 c&d).

Next to accurate information on taxonomic composition and taxon richness, recognition of closely related strains is required from amplicon sequence processing tools. Six bacterial genera were represented by two strains each in the bacterial dataset and recognized as such by ASVs. In the case of three prokaryotic genera, the true diversity was not resolved by ASVs, with three *Thermotoga* strains and two *Salinispora* and two *Sulfitobacter* strains conflated as two and one strains, respectively (Supplementary table 3). Micro-diversity was correctly identified for two strains of *Aspergillus* and the three *Fusarium strains* (although one was misclassified) for the fungal dataset. Strain-diversity was overestimated for the fungal dataset in *Rhizophagus irregularis,* which is known to contain within-genome diversity of ribosomal RNA gene sequences [44]. Overall, dadasnake returns accurate results for taxonomic composition, richness and micro-scale diversity within the limits of taxonomic resolution within short regions.

### Use cases: limitations

The analysis of the mock community data also revealed limitations of the approach in general. A commonly used approach to detect underestimation of richness at low sequencing depths is to plot rarefaction curves or use richness estimators [45–47], which use subsamples of the assigned reads to model how much the addition of further sequencing would increase the observed richness. However, the statistical requirements for delineation of ASVs mean that not all sequenced taxa are represented by an ASV in a given data set (https://github.com/benjjneb/dada2/issues/317). This in turn leads to the flattening of rarefaction curves derived from finished ASV tables, although an increase in real sequencing depth would lead to a greater number of observed ASVs (Figure 3 c&d). Richness estimates and rarefaction curves based on DADA2 datasets need to be handled with caution and whenever richness estimates are essential should be based on subsamples that are processed by DADA2 independently rather than post-hoc models.

**Figure 3:**
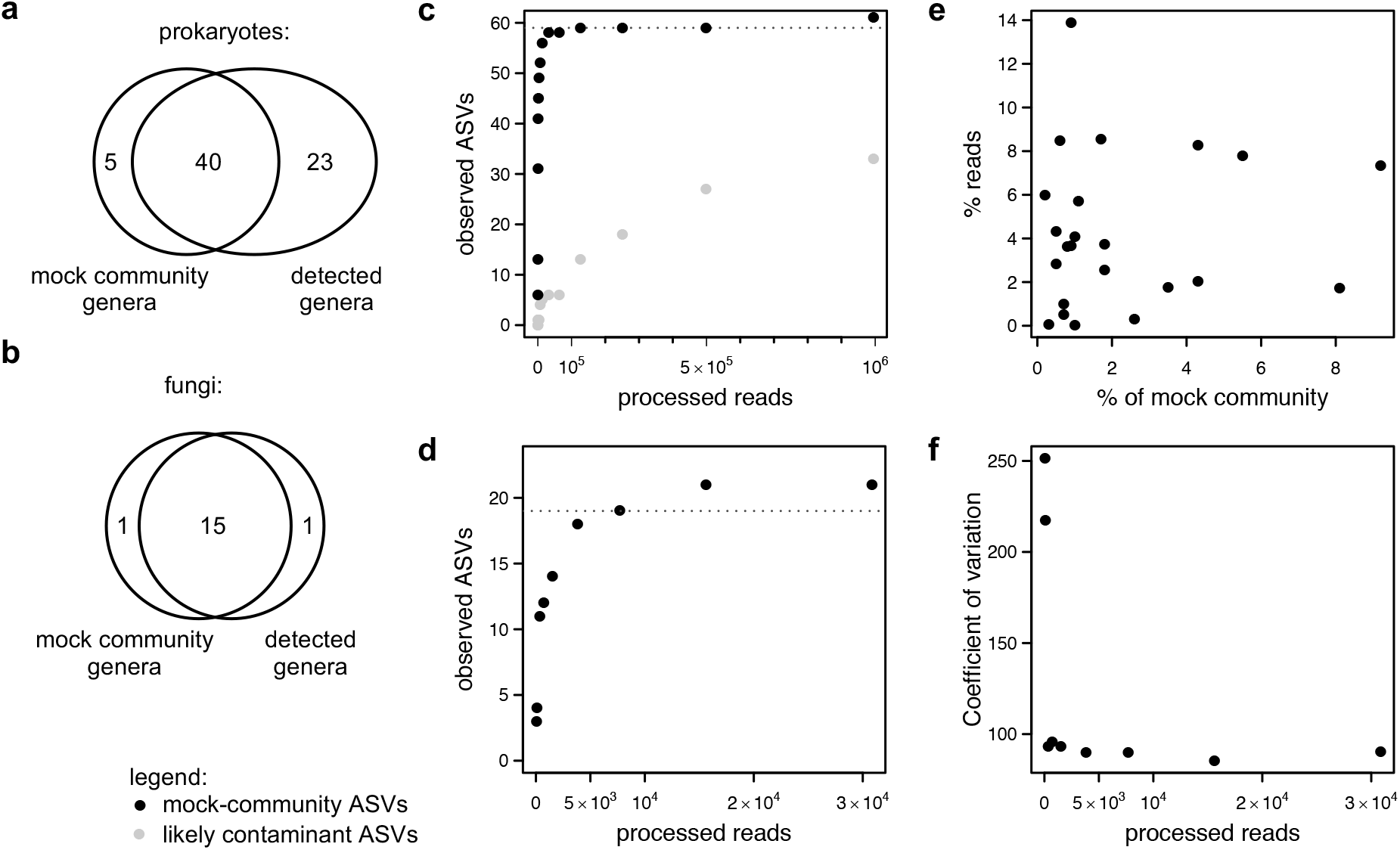
Comparison of mock community composition with analysis results. a) Detection of prokaryotic genera at the highest sequencing depth (1.6 mio reads); b) detection of fungal genera at the highest sequencing depth (40,000 reads); c) number of detected prokaryotic ASVs plotted against the number of processed (non-chimeric) reads - black circles: ASVs of taxa from the mock community, grey circles: likely contaminant taxa; d) number of detected fungal ASVs against the number of processed (non-chimeric) reads of the fungal mock community; c & d) dotted lines indicate expected taxa richness; e) missing correlation of real percentages of the mock communities and detected relative abundances of prokaryotic genera; f) coefficients of variation between relative abundances of taxa that should be equally abundant in the fungal mock community.

A second limitation, common to amplicon sequencing, is that relative abundances of ASVs are not reflective of the actual abundance of the sequenced taxa, which varied for the prokaryotic mock community and were equal in the fungal mock community. Specifically, the relative abundance of the prokaryotic taxa did not correlate with the relative abundance of reads (Figure 2 e). The relative abundance of reads for the fungal taxa varied by several orders of magnitude, despite equal inputs (Figure 3 f). There are numerous reasons for misrepresentation of abundances by PCR-based analyses [48]. Of note, the variation in the relative abundance estimates is observed to be highest at low sequencing depths (Figure 3 e&f). Therefore, whenever comparisons of relative abundances within samples are undertaken, it is necessary to, at the least, ensure that sequencing depths of all samples are sufficient to reach stable estimates. However, the analysis of the mock community case studies also suggests that true relative abundances can never be determined, which should be accounted for in experimental design and interpretation.

## Methods

### Bacterial and archaean mock community dataset

The largest library of the Illumina sequencing datasets of a 59 species mock-community [49], comprising 10 archaea and 49 bacteria (composition see Supplementary table 3) was retrieved from the European Nucleotide Archive ENA under accession ERR777696. The ground-truth composition of the mock-community was manually extracted from the publication and the taxonomic names adapted to the convention of the SILVA v. 138 database [50]. To analyse the effect of sequencing depth on the recovery of the mock-community, the dataset was subsampled to 100, 200, 500, 1,000, 2,000, 5,000, 10,000, 20,000, 50,000, 100,000, 200,000, 400,000, 800,000, and 1,600,000 read pairs.

The same configuration was used to run dadasnake on all subsamples. The most important settings include removal of the primers from either read (515F, specified as 5-GTGYCAGCMGCCGCGGTAA, and 806R, specified as 5-GGACTACNVGGGTWTCTAAT, with a maximum of 20 % mismatch); truncation of the reads at positions with a quality below 13, before removal of forward and reverse reads with less than 170 and 130 nt length, respectively, and truncation to these lengths before removal of reads with an expected error above 0.2; a minimum of 12 bp overlap was required for merging of denoised sequences; chimeras were removed on consensus.

### Fungal mock community sequencing

The ITS2 region of an even 19 species fungal mock community [42] provided by Matt Bakker (composition see Supplementary table 3), was amplified using the primers F-ITS4 5-TCCTCCGCTTATTGATATGC [51] and R-fITS7 5-GTGARTCATCGAATCTTTG [52] modified with heterogeneity spacers according to [53]. Amplicon libraries were prepared using the Nextera XT kit (Illumina) and sequenced on an Illumina MiSeq with v.3 chemistry at 2 x 300 bp. Sequencing was performed in triplicates and all reads were pooled for the analysis presented here. The sequencing data is accessible at the NCBI Short Read Archive under BioProject accession PRJNA626434. The groundtruth composition of the data was manually extracted from the publication and the taxonomic names were adjusted to the ones used in the Unite 8.0 database. To analyse the effect of sequencing depth on the recovers of the mock-community, the dataset was subsampled to 100, 200, 500, 1,000, 2,000, 5,000, 10,000, 20,000 and 40,000 reads.

The same configuration was used for running dadasnake on all subsamples. The most important settings were: removal of the primers from either read with a maximum of 20 % mismatch; truncation of the reads at positions with a quality below 15, before removal of reads with less than 70 nt length and removal of reads with an expected error above 3; a minimum of 20 bp overlap was required for merging of denoised sequences; chimeras were removed on consensus; ITSx was run on the ASVs which would remove non-fungal ASVs (which did not occur in the mock-community).

### Performance testing

To demonstrate dadasnake’s performance on a small laptop computer, a small data set of 24 16S rRNA gene amplicon sequences from a local soil fertilization study [39] were downloaded from the NCBI short read archive (PRJNA517390) using the fastq-dump function of the SRA-toolkit. Using the settings optimized for the bacterial mock-community, dadasnake was run either on a computer cluster using up to 1 or 4 threads with 8 GB RAM each, or without cluster-mode on three cores of a laptop with an Intel i5-2520M CPU with 2.5 GHz and 8 GB shared RAM. The same settings as for the mockcommunity with bacteria and archea was used.

To compare performance of dadasnake on a medium sized study in different settings, ITS1 amplicon sequences of 267 samples measured using Illumina HiSeq technology in a global study on fertilization effects [40] were downloaded from the NCBI short read archive (PRJNA272747) using the fastq-dump function of the SRA-toolkit. Owing to the variable length of the ITS1 region, reads were not truncated to a specified length, but trimmed to a minimum per-base quality of 15 (also discarding reads with a higher maximum expected error than 3). After error modelling and ASV construction per sample, read pairs were merged with at least 20 bp overlap, allowing for 2 mismatches. After table setup, the ITSx classifier was run to remove non-fungal ASVs before taxonomic annotation (using the mothur [14] classifier). The same runs were performed on either a compute cluster using up to 50 threads or only up to 4 threads with 8GB RAM each.

27,081 samples analysed by the Earth Microbiome Project [12] stored under accessions ERP021896, ERP020023, ERP020508, ERP017166, ERP020507, ERP017221, ERP016412, ERP020884, ERP020022, ERP020510, ERP017438, ERP016395, ERP020539, ERP016468, ERP020590, ERP020021, ERP020587, ERP020560, ERP020589, ERP017176, ERP017220, ERP017174, ERP016405, ERP020591, ERP021691, ERP016416, ERP022167, ERP021699, ERP016495, ERP022245, ERP016748, ERP016749, ERP016752, ERP016540, ERP006348, ERP016543, ERP016746, ERP016586, ERP016735, ERP021864, ERP016588, ERP016587, ERP016539, ERP016734, ERP016492, ERP003782, ERP016607, ERP016581, ERP016557, ERP016464, ERP016542, ERP016541, ERP016591, ERP016854, ERP016852, ERP016286, ERP016451, ERP023684, ERP016869, ERP010098, ERP016879, ERP016883, ERP016466, ERP016496, ERP016880, ERP016455, ERP016900, ERP016924, ERP016923, ERP016925, ERP016927, ERP016469, ERP016329, ERP016926, ERP021540, ERP021541, ERP021542, ERP021543, ERP021544, ERP021545, ERP016937, ERP016131, ERP016483, ERP016252, ERP022166, ERP016414, ERP016472, ERP023686, ERP017459, ERP016287, ERP016285, ERP005806, ERP021895, ERP016384, ERP016491, and ERP006348 were downloaded from the NCBI short read archive using the fastq-dump function of the SRA-toolkit. In accordance with the published analysis reads were trimmed to 90 bp, before quality control (discarding reads with a higher maximum expected error than 0.2 or positions with less than 13 quality score), error modelling (per project accession), ASV construction (per sample), table set-up, and taxonomic annotation (using the mothur [14] classifier; configuration see Supplementary file 1). To handle the combined dataset table, 360 GB RAM were reserved for the final steps in R.

### Databases

The SILVA [50] RefSSU_NR99 database v. 138 was used for the taxonomic classification of bacterial and archaean ASVs. Fungal ASVs were classified against the UNITE v8 database [54,55]. Both sets of ASVs were classified using the Bayesian classifier as implemented in mothur’s classify.seqs command [14], with a cut-off of 60.

### Visualization and statistics

The output of all dadasnake runs of mock communities was gathered in an R-workspace (tabular version see Supplementary table 3). The coefficient of variation was calculated as the ratio of the standard deviation to the mean. The cluster-job information for the performance tests was gathered in an R-workspace. Efficiency was calculated as the ratio of CPU-time divided by the product of used slots and real walltime.

## Availability of supporting source code and requirements

Project name: dadasnake

Project home page: https://github.com/a-h-b/dadasnake

Operating system(s): Linux

Programming language: Python, R, bash

Other requirements: anaconda or other conda package manager

License: GNU GPL-3.0

## Availability of supporting data

The raw sequencing data generated for this manuscript are accessible on NCBI’s Sequence Read Archive under BioProject accession PRJNA626434. Processing results of the mock community data sets, the ground-truth mock community compositions, and the scripts to visualize the use case datasets are available from Zenodo https://doi.org/10.5281/zenodo.4104624.

## List of abbreviations

OTU: operational taxonomic unit
ASV: amplicon sequence variants (=ESV)
ESV: exact sequence variants (=ASV)
rRNA: ribosomal RNA

## Authors’ contributions

Conceptualization, software, analysis, writing: AH-B; optimization and testing: CW; sequencing: BS. All authors contributed to the manuscript text and approve its contents.

## Acknowledgements

The authors would like to acknowledge Kezia Goldmann and Julia Moll for testing early versions of the workflow; François Buscot for funding acquisition and providing resources; Guillaume Lentendu for discussions. Data processing has been performed at the High-Performance Computing (HPC) Cluster EVE, a joint effort of both the Helmholtz Centre for Environmental Research - UFZ and the German Centre for Integrative Biodiversity Research (iDiv) Halle-Jena-Leipzig and the authors thank Christian Krause and the other administrators for excellent support. Matthew Bakker is acknowledged for the generous provision of the fungal mock community.

